# Evolutionary rates are correlated between cockroach symbiont and mitochondrial genomes

**DOI:** 10.1101/542241

**Authors:** Daej A. Arab, Thomas Bourguignon, Zongqing Wang, Simon Y. W. Ho, Nathan Lo

**Author notes:** Authors for correspondence: Daej A. Arab, Nathan Lo.

## Abstract

Bacterial endosymbionts evolve under strong host-driven selection. Factors influencing host evolution might affect symbionts in similar ways, potentially leading to correlations between the molecular evolutionary rates of hosts and symbionts. Although there is evidence of rate correlations between mitochondrial and nuclear genes, similar investigations of hosts and symbionts are lacking. Here we demonstrate a correlation in molecular rates between the genomes of an endosymbiont (*Blattabacterium cuenoti*) and the mitochondrial genomes of their hosts (cockroaches). We used partial genome data for multiple strains of *B. cuenoti* to compare phylogenetic relationships and evolutionary rates for 55 cockroach/symbiont pairs. The phylogenies inferred for *B. cuenoti* and the mitochondrial genomes of their hosts were largely congruent, as expected from their identical maternal and cytoplasmic mode of inheritance. We found a correlation between evolutionary rates of the two genomes, based on comparisons of root-to-tip distances and on comparisons of the branch lengths of phylogenetically independent species pairs. Our results underscore the profound effects that long-term symbiosis can have on the biology of each symbiotic partner.

## 1. Introduction

Rates of molecular evolution are governed by a multitude of factors and vary significantly among species [1, 2]. In the case of symbiotic organisms, such rates may be influenced by the biology of their symbiotic partner, in addition to their own. This is particularly the case for strictly vertically transmitted, obligate intracellular symbionts (hereafter ‘symbionts’), which have a highly intimate relationship with their hosts [3]. For example, a small host effective population size will potentially lead to increased fixation of slightly deleterious mutations within both host and symbiont genomes, owing to the reduced efficacy of selection.

When the phylogenies of host and symbiont taxa are compared, simultaneous changes in evolutionary rate between host-symbiont pairs might be evident in their branch lengths. Some studies have found a correlation in evolutionary rates between nuclear and mitochondrial genes in sharks [4], herons [5], and turtles [6], suggesting that host biology affects substitution rates in nuclear and cytoplasmic genomes in similar ways. In insects, nuclear genes that interact directly with mitochondrial proteins have shown rate correlations with mitochondrial genes [7].

Potential correlations in evolutionary rates between hosts and bacterial symbionts remain untested. Evidence for correlated levels of synonymous substitutions was found in a study of one nuclear gene and two mitochondrial genes from *Camponotus* ants and three genes from their *Blochmannia* symbionts [8]. However, the study did not determine whether this correlation was driven by rates of evolution, time since divergence, or both. Numbers of substitutions tend to be low for closely related pairs of hosts and their corresponding symbionts, and high for more divergent pairs, leading to a correlation with time that does not necessarily reflect correlation in evolutionary rates.

*Blattabacterium cuenoti* (hereafter *Blattabacterium*) is an intracellular bacterial symbiont that has been in an obligatory intracellular and mutualistic relationship with cockroaches for over 200 million years [9, 10]. Found in highly specialized cells in the fat bodies of cockroaches, *Blattabacterium* is required for host fitness and fertility, and is transovarially transmitted from the mother to the progeny [11, 12]. Genome-wide analyses of the symbiont have confirmed its role in host nitrogen metabolism and the synthesis of essential amino acids [13, 14]. The genomes of 21 *Blattabacterium* strains sequenced to date are highly reduced compared with those of their free-living relatives, ranging in size from 590 to 645 kb [15, 16]. They contain genes encoding enzymes for DNA replication and repair, with some exceptions (*holA*, *holB*, and *mutH*) [13, 16, 17]. The extent to which host nuclear proteins are involved in the cell biology of *Blattabacterium*, and particularly DNA replication, is not well understood.

We recently performed a study of cockroach evolution and biogeography using mitochondrial genomes [9]. During this process, we obtained partial genomic information for several *Blattabacterium* strains. These data provide the opportunity to test for correlation of molecular evolutionary rates between *Blattabacterium* and host-cockroach mitochondrial DNA. Here we infer phylogenetic trees for 55 *Blattabacterium* strains on the basis of 104 genes and compare branch lengths and rates of evolution for host-symbiont pairs across the phylogeny. We find evidence of markedly increased rates of evolution in some *Blattabacterium* lineages, which appear to be matched by increased rates of evolution in mitochondrial DNA of host lineages.

## 2. Materials and methods

A list of samples and collection data for each cockroach examined is provided in table S1 (see electronic supplementary material, ESM). For the majority of taxa examined in this study, we obtained *Blattabacterium* sequence data from genomic libraries originally used in a previous study of cockroach mitochondrial genomes carried out by our laboratories [9]. In some cases, new genomic data were obtained from fat bodies of individual cockroaches (see ESM for further details). We obtained 104 genes of 55 *Blattabacterium* strains from these data and aligned them with orthologues from seven outgroup taxa from Flavobacteriales (details provided in ESM).

Genomic data were assembled and annotated, and then aligned and tested for saturation. After the exclusion of 3rd codon sites in each data set, total lengths for the mitochondrial and *Blattabacterium* alignments were 11,051 bp and 71,458, respectively. The former was partitioned into four subsets (1st codon sites, 2nd codon sites, rRNAs, and tRNAs), and the latter into two subsets according to codon positions. Trees were inferred using maximum likelihood in RAxML v8.2 [18], using 1000 bootstrap replicates to estimate node support. We examined congruence between host and symbiont phylogenies using the distance-based ParaFit [19] in R 3.5.1 [20]. Root-to-tip distances from the RAxML analyses for each host and symbiont pair were subjected to Pearson correlation analysis. Branch-length differences between hosts and symbionts were compared for 27 phylogenetically independent species pairs across the topology (see figure S1 in ESM). These were calculated using a fixed topology (derived from the *Blattabacterium* analysis described above) for each of the following three data sets: 1) 1st+2nd codon sites of protein-coding genes; 2) translated amino acid sequences; 3) 1st+2nd codon positions of protein-coding genes plus the inclusion of rRNAs and tRNAs in the case of the mitochondrial data set. Further details on phylogenetic methods are provided in the ESM.

## 3. Results

In all analyses, there was strong support for the monophyly of each cockroach family with the exception of Ectobiidae (figure 1). The topologies inferred from the host and symbiont data sets were congruent (*p* = 0.001). In only two cases was a disagreement found to be supported by >85% bootstrap support in both trees (the sister group of Lamproblattidae+Anaplectidae; the sister group to *Carbrunneria paraxami*+*Beybienkoa kurandensis*).

**Figure 1.**
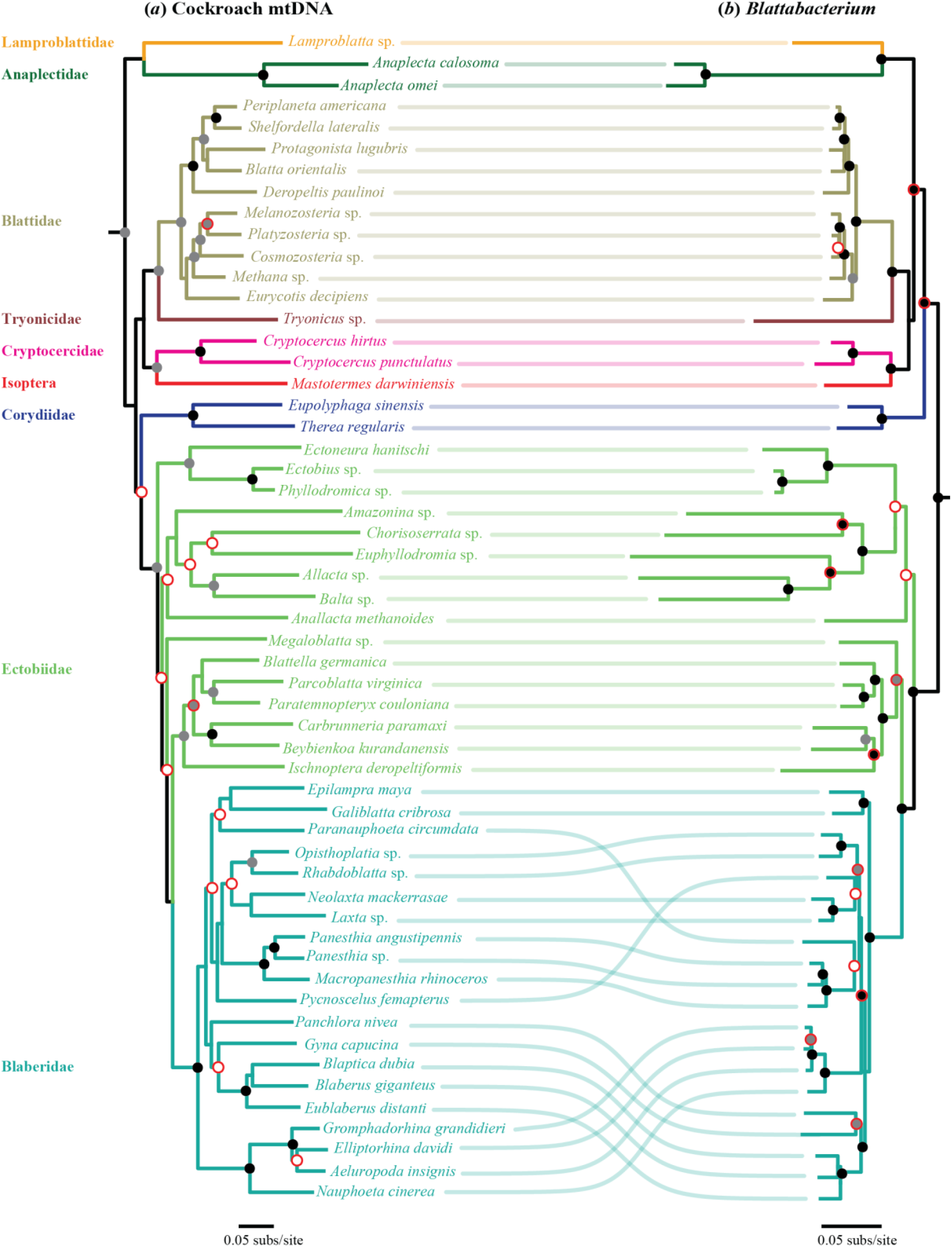
Congruence between (*a*) phylogenetic tree of host cockroaches inferred using maximum likelihood from whole mitochondrial genomes, and (*b*) phylogenetic tree of *Blattabacterium* inferred using maximum likelihood from 104 protein-coding genes (3rd codon sites excluded from both data sets). Shaded circles at nodes indicate bootstrap values (black = 100%, grey = 85–99%). Nodes without black or grey circles have bootstrap values <85%. Red outlines on circles indicate disagreement between the phylogenies. Branches are coloured according to their membership of different cockroach families.

We found a correlation between root-to-tip distances for protein-coding genes from hosts and their symbionts (*R* = 0.75, figure 2*a*). Similar results were found when rRNAs and tRNAs were included in the host data set (*R* = 0.73, figure S4*a*). The highest rates of evolution in the host and symbiont data sets (on the basis of branch lengths; see figure 1) were in members of an ectobiid clade containing *Allacta* sp., *Amazonina* sp., *Balta* sp., *Chorisoserrata* sp., and *Euphyllodromia* sp., and a separate clade containing the Anaplectidae. After excluding these taxa, evolutionary rates remained correlated, although to a lesser degree (*R* = 0.35, figure 2*b*). The sharing of branches between taxa in the estimation of root-to-tip distances renders the data in these plots phylogenetically non-independent and precludes statistical analysis.

**Figure 2.**
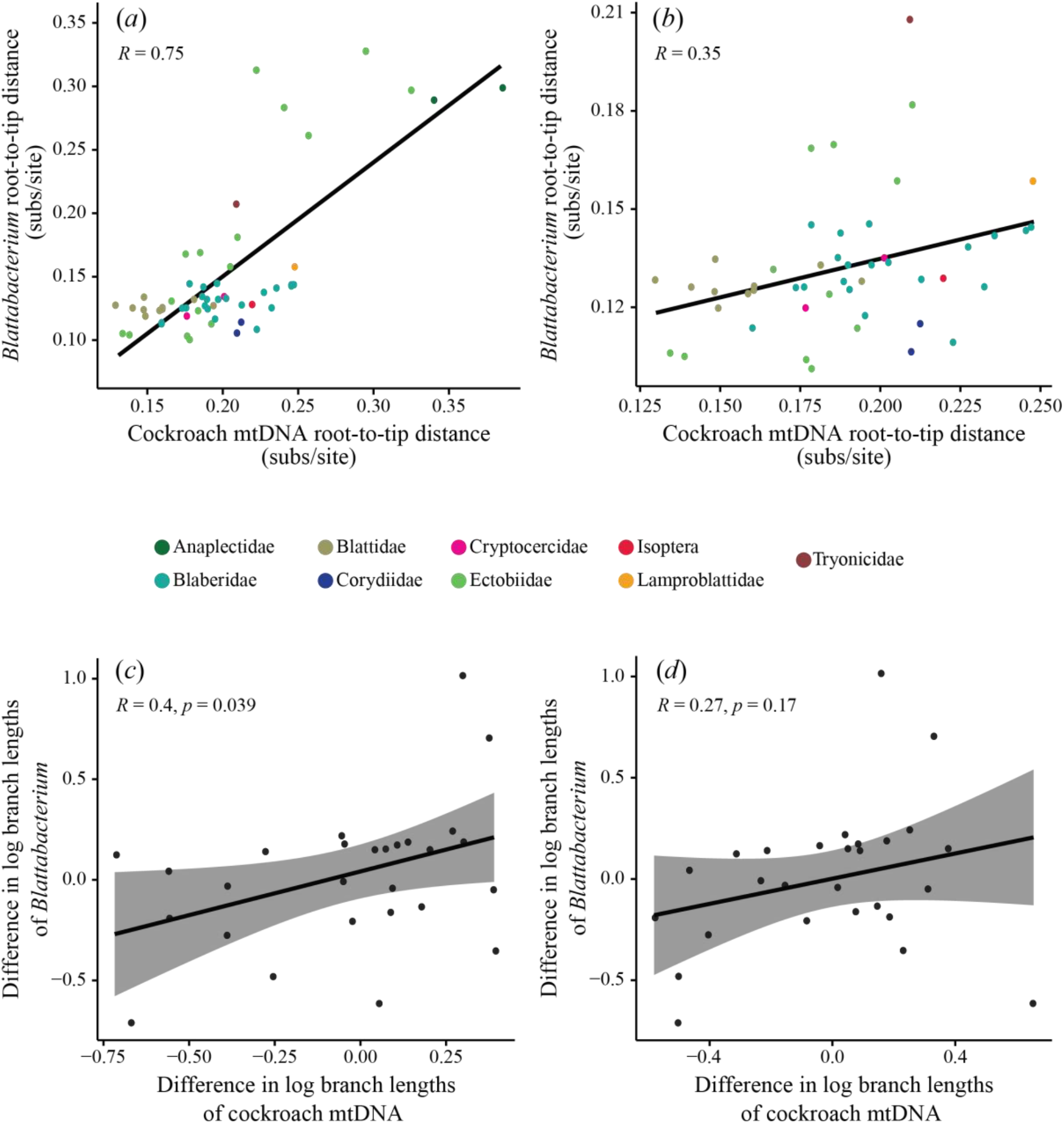
Comparison of evolutionary rates of *Blattabacterium* symbionts and their host cockroaches. (*a*) Correlation of root-to-tip distances in phylogenies of *Blattabacterium* and cockroaches, inferred using maximum-likelihood analysis of protein-coding genes from each data set, with 3rd codon sites excluded. (*b*) Correlation of root-to-tip differences following the removal of five rapidly evolving ectobiid taxa (*Amazonina* sp., *Chorisoserrata* sp., *Allacta* sp., *Balta* sp., and *Euphyllodromia* sp.) and two anaplectids. Colours represent data from representatives of different cockroach families, as shown in the colour key. Grey circles represent internal branches. (*c*) Correlation of log-transformed branch-length differences between phylogenetically independent pairs of host and symbiont taxa, based on protein-coding genes only, and (*d*) with the addition of rRNAs and tRNAs to the host mitochondrial data set.

A comparison of branch lengths among phylogenetically independent pairs of host and symbiont taxa based on protein-coding genes revealed a significant correlation between their rates of evolution (*R* = 0.40, *p* = 0.039; figure 2*c*, figure S1). Equivalent analyses of branch lengths inferred from amino acid data also revealed a significant rate correlation between host and symbiont (*R* = 0.43, *p* = 0.023; figure S3*a*). However, there was no rate correlation between host and symbiont following inclusion of rRNAs and tRNAs in the host mitochondrial data set (*R* = 0.27, *p* = 0.17; figure 2*d*). Analyses involving standardization of branch-length differences yielded significant rate correlations for the protein-coding gene and amino acid data sets (*R* = 0.43–0.48, *p* = 0.011–0.023; see figures S2, S3), and mixed results in the case of the inclusion of rRNAs and tRNAs in the host mitochondrial data set (*R* = 0.34–0.40, *p* = 0.041–0.085; see figure S4).

## 4. Discussion

We have detected a correlation in molecular evolutionary rates between *Blattabacterium* and host mitochondrial genomes, using two different methods of analysis. To our knowledge, this is the first evidence of such a correlation in a host-symbiont relationship. Previous studies found a correlation in evolutionary rates between mitochondrial and nuclear genes in various animal groups [4–7].

Similar forces acting on the underlying mutation rates of both host and symbiont genomes could translate into a relationship between their substitution rates. This could potentially occur if symbiont DNA replication depends on the host’s DNA replication and repair machinery [21]. A better understanding of the level of integration of host-encoded proteins in the metabolism of *Blattabacterium* will allow further exploration of this issue. A study of insect genomic data found a correlation in rates between mitochondrial genes and nuclear genes that encode proteins targeted to mitochondria [7]. However, since *Blattabacterium* is not known to be directly associated with mitochondria, interactions between symbiont and organellar proteins are unlikely to explain the correlations in rates that we have found here.

Short host generation times could potentially lead to elevated evolutionary rates in host and symbiont [22], assuming that increased rates of symbiont replication are associated with host reproduction, as is found in *Blochmannia* symbionts of ants [23]. Variations in metabolic rate and effective population size between host taxa could also explain the rate correlations that we have observed. Unfortunately, with the exception of a few pest and other species, generation time, metabolic rates, and effective population sizes are poorly understood in cockroaches. This precludes an examination of their influence on evolutionary rates in host and symbiont.

The addition of mitochondrial rRNA and tRNA data weakened the correlations found in the branch-length comparisons of species pairs. The reasons for this are unclear but they might be associated with the conserved nature of tRNAs and the stem regions of rRNAs, or highly variable loop regions in the latter.

*Blattabacterium* is a vertically transmitted, obligate intracellular mutualistic symbiont, whose phylogeny is expected to mirror that of its hosts. This is especially the case for phylogenies inferred from mitochondrial DNA, since mitochondria are linked with *Blattabacterium* through vertical transfer to offspring through the egg cytoplasm. As has been found in previous studies [24–26], we observed a high level of agreement between the topologies inferred from cockroach mitochondrial and *Blattabacterium* genome data sets. In some cases, however, disagreements were observed between well-supported relationships. The variability in rates that we observed between some lineages, and/or the highly increased rate of mitochondrial DNA compared with *Blattabacterium* DNA, could be responsible for these disagreements. Owing to long periods of co-evolution and co-cladogenesis between cockroaches and *Blattabacterium* [9, 24], potential movement of strains between hosts (for example, via parasitoids) is not expected to result in the establishment of new symbioses, especially between hosts that diverged millions of years ago.

In conclusion, our results highlight the profound effects that long-term symbiosis can have on the biology of each symbiotic partner. The rate of evolution is a fundamental characteristic of any species; our study indicates that it can become closely linked between organisms as a result of symbiosis. Further studies are required to determine whether the correlation that we have found here also applies to the nuclear genome of the host. Future investigations of generation time, metabolic rate, and effective population sizes in cockroaches and *Blattabacterium* will allow testing of their potential influence on evolutionary rates.

## Supporting information

Electronic supplementary material

## Authors’ contributions

N.L., D.A.A., T.B., and S.Y.W.H. designed the study. Z.W. and T.B. collected and provided specimens; D.A.A., T.B. generated sequence data; D.A.A., T.B, N.L., and S.Y.W.H. performed data analysis and interpreted results; N.L. and D.A.A. wrote the manuscript, with contributions from T.B. and S.Y.W.H. All authors approved the final version of the manuscript and agree to be accountable for all aspects of the work.

## Ethics

This article does not present research with ethical considerations.

## Data accessibility

Sequence data have been uploaded to GenBank (accession numbers MN038417–MN043259) and alignments have been uploaded to DRYAD: DOI: https://doi.org/10.5061/dryad.v6wwpzgqw.

## Funding

D.A.A. was supported by an International Postgraduate Research Stipend from the Australian Government. S.Y.W.H. and N.L. were supported by Future Fellowships from the Australian Research Council. T.B. was supported by the Japan Society for the Promotion of Science KAKENHI 18K14767, and by an EVA4.0 grant (No. CZ.02.1.01/0.0/0.0/16_019/0000803) from the OP RDE. Z.W. acknowledges funding from the National Natural Sciences Foundation of China (31872271).

## Competing interests

There are no competing interests.

## Acknowledgements

We thank Qian Tang, Frantisek Juna, James Walker, and David Rentz for providing specimens, and Charles Foster for assistance with data curation.

## Notes

#### Summary of Updates

Ethics statement updated. Data accessibility updated. Figure 2 higher resolution. Modified Methods and results sections Modified Figures 1 and 2

