## Supplementary material for "Evolutionary rates are correlated between cockroach symbiont and mitochondrial genomes": Electronic supplementary material

### Sampling and *Blattabacterium* genomic data

A list of samples and collection data for each cockroach and outgroup examined is provided in table S1. All specimens examined in this study are stored at the Okinawa Institute of Science and Technology, Japan. For the majority of taxa examined in this study, we obtained *Blattabacterium* sequence data from genomic libraries originally used in a previous study of cockroach mitochondrial genomes carried out by our laboratories [1]. In some cases, new genomic data were obtained from fat bodies of individual cockroaches, as follows. DNA was extracted using a DNeasy Blood and Tissue kit (Qiagen), according to the manufacturer's protocol. Individual DNA samples were tagged with unique barcode combinations, mixed in equimolar concentration, and 150 bp paired-end-reads-sequenced with an Illumina HiSeq4000, following the methods described previously [1].

For each cockroach species, raw sequence data from the previous study [1] or the current study were assembled using CLC, and subject to blastn analysis using the published *Blattabacterium* genomes from *Blattella germanica* [2], *Periplaneta americana* [3], and *Cryptocercus punctulatus* [4] as subject sequences. Contigs identified as being derived from *Blattabacterium* during this step were then annotated using Prokka v1.12 [5]. Across the 55 strains examined in this study, a total of 104 orthologous genes were used for analysis. These were found in ≥95% of all taxa. All taxa had over 90% of 104 genes, except for *Aeluropoda insignis* which only had 83 (79.8%) genes. Missing genes were presumed to be a result of uneven sequencing coverage of samples and the relatively low sequencing coverage used, rather than the actual absence of these genes from their genomes; further work is required to confirm their presence or absence. The genome sequences of outgroups were obtained from GenBank and included three strains of *Sulcia muelleri* (accession numbers CP002163, AP013293, and CP010828), one *Flavobacterium gilvum* (CP017479), one *Lutibacter* sp. (CP017478), one *Tenacibaculum dicentrarchi* (CP013671), and one *Polaribacter* sp. (LT629752).

The 104 orthologous *Blattabacterium* genes were each aligned at the amino acid level individually using TranslatorX [6] and concatenated into a 107,187 bp alignment. The mitochondrial genome data set included all protein-coding genes from each taxon plus 12S rRNA, 16S rRNA, and the 22 tRNA genes. Mitochondrial protein-coding genes were aligned using TranslatorX, while MAFFT [7] was used to align 12S rRNA, 16S rRNA, and the 22 tRNAs. All mitochondrial sequences were then concatenated into a 14,802 bp alignment. We tested for substitution saturation using Xia's method implemented in DAMBE 6 [8, 9]. Third codon sites in the mitochondrial data set were saturated (NumOTU = 32, $I_{ss}$ = 0.804, $I_{ss.cAsym}$ = 0.809) and were excluded from our analyses. Although the *Blattabacterium* sequences were not significantly saturated at 3rd codon sites (NumOTU = 32, $I_{ss}$ = 0.649, $I_{ss.cAsym}$ = 0.819), we excluded these sites from our analyses because the test statistic was close to the critical value. After the exclusion of 3rd codon sites, the total lengths of the final data sets were 11,051 bp and 71,458 for the mitochondrial and *Blattabacterium* alignments, respectively.

**Phylogenetic analysis**

Maximum-likelihood phylogenetic analyses were carried out in RAxML v8.2 [10], using 1000 bootstrap replicates to estimate node support. The cockroach mtDNA data set was partitioned into four subsets: 1st codon sites, 2nd codon sites, rRNAs, and tRNAs. The *Blattabacterium* data set was partitioned into two subsets: 1st codon sites and 2nd codon sites. For each data set, a separate GTR+G substitution model was assigned to each subset.

We used ParaFit in R 3.5.1 [11] to quantify congruence between host and symbiont topologies. We first created matrices of patristic distances calculated from maximum-likelihood host and symbiont phylogenies and a host-symbiont association matrix. We then performed a global test with 999 permutations, using the ParafitGlobal value and a *p*-value threshold of 0.05 to determine significance.

**Root-to-tip distances and comparison of phylogenetically independent pairs of host and symbiont branch lengths**

Root-to-tip distances from the RAxML analyses for each host and symbiont pair were calculated and subjected to Pearson correlation analysis using the R packages ape [12], phylobase [13], and adephylo [14]. The use of root-to-tip distances removes the confounding effects of time, because all lineages leading to the tips of the tree have experienced the same amount of time since evolving from their common ancestor. However, the sharing of internal branches by groups of taxa renders these data non-independent. Therefore, we compared branch-length differences between hosts and symbionts for 27 phylogenetically independent species pairs across the topology (see figure S1). These were calculated using a fixed topology (derived from the *Blattabacterium* analysis described above) for each of the following three data sets: 1) 1st+2nd codon sites of protein-coding genes; 2) translated amino acid sequences; and 3) 1st+2nd codon positions of protein-coding genes plus the inclusion of rRNAs+tRNAs in the case of the mitochondrial data set. Branch lengths were log transformed, and differences between pairs of hosts and pairs of symbionts were calculated and compared via Pearson correlation analysis.

To test for potential biases in the data that violate the assumptions of linear regressions, we compared the absolute mean value of log-transformed branch lengths with the log-transformed branch-length differences [15]. We found no significant correlation between these values ($R$ = 0.07, $p$ = 0.63 for data from host cockroaches; $R$ = 0.06, $p$ = 0.66 for data from *Blattabacterium*), indicating that the data were suitable for use in our analyses. We also performed analyses in which branch-length differences were standardized following previous recommendations [16], to account for the potential confounding effects of the different amounts of time that sister pairs have had to diverge. Three standardizations were carried out, each based on dividing log-transformed branch-length differences by the square root of an estimate of time since divergence for the pair. In the first, time since divergence for host pairs was estimated as the average branch length of the host pair, divided by an assumed rate of 0.001 subs/site/million years, while for corresponding symbionts it was estimated as the average branch length of the symbiont pair, divided by the same assumed rate. In the second and third standardizations, times since divergence for both symbionts and hosts were based either on average branch lengths of host pairs only or symbiont pairs only.

**Supplementary table S1.** A list of samples and collection data for each cockroach examined.

| Species | Family | Sample ID | Collecting locality | Collector | Date |
|---|---|---|---|---|---|
| ***Aeluropoda insignis*** | Blaberidae | B002 | Breeding colony of Kyle Kandilian | N/A | N/A |
| ***Allacta australiensis*** | Ectobiidae | AUS Allacta | James Cook University, Rainforest site, Queensland, Australia | David Rentz | 22-Jun-2015 |
| ***Amazonina* sp.** | Ectobiidae | Z256E | Ecuador, Bosque Protector del Alto Nangaritza | Frantisek Juna | Apr-2016 |
| ***Anallacta methanoides*** | Ectobiidae | B057 | Breeding colony of Kyle Kandilian | N/A | N/A |
| ***Anaplecta calosoma*** | Anaplectidae | Cockroach contig 1688 | Kuranda, Queensland, Australia | David Rentz | 17-Nov-2015 |
| ***Anaplecta omei*** | Anaplectidae | Anaplecta_omei | Mt Emei, Sichuan, China | Zongqing Wang | 01-Jul-2013 |
| ***Balta* sp.** | Ectobiidae | Balta_sp. | Cairns, Queensland, Australia | David Rentz | 18-Dec-2015 |
| ***Beybienkoa kurandanensis*** | Ectobiidae | Beybienkoa_kurandanensis | Cairns, Queensland, Australia | David Rentz | 18-Dec-2015 |
| ***Blaberus giganteus*** | Blaberidae | BGIGA | GenBank | N/A | N/A |
| ***Blaptica dubia*** | Blaberidae | B056 | Breeding colony of Kyle Kandilian | N/A | N/A |
| ***Blatta orientalis*** | Blattidae | BOR | GenBank | N/A | N/A |
| ***Blattella germanica*** | Ectobiidae | BGE | GenBank | N/A | N/A |
| ***Carbrunneria paramaxi*** | Ectobiidae | Carbru | Cairns, Queensland, Australia | David Rentz | 05-Oct-2015 |
| ***Chorisoserrata* sp.** | Ectobiidae | CHORI | Yunnan, China | Zongqing Wang | 01-Jul-2013 |
| ***Cosmozosteria* sp.** | Blattidae | B117 | Cape Upstart, Queensland, Australia | James Walker | 13-Oct-2015 |
| ***Cryptocercus hirtus*** | Cryptocercidae | HIR | Mt Taibai, Shaanxi, China | N/A | 06-Oct-2013 |
| ***Cryptocercus punctulatus*** | Cryptocercidae | CPU | GenBank | N/A | N/A |
| ***Deropeltis paulinoi*** | Blattidae | B069 | Breeding colony of Kyle Kandilian | N/A | N/A |
| ***Ectobius* sp.** | Ectobiidae | Z254C | Slovenia | Frantisek Juna | Apr-2016 |
| ***Ectoneura hanitschi*** | Ectobiidae | Ectoneura_hanitschi | James Cook University, Rainforest site, Queensland, Australia | David Rentz | 18-Dec-2015 |
| ***Epilampra maya*** | Blaberidae | B095 | Arcadia, Florida, USA | Kyle Kandilian | 07-Jul-2009 |
| ***Eublaberus distanti*** | Blaberidae | B025 | Breeding colony of Kyle Kandilian | N/A | |
| ***Euphyllodromia* sp.** | Ectobiidae | Z257 | Podocarpus National Park, Ecuador | Frantisek Juna | Apr-2016 |
| ***Eupolyphaga sinensis*** | Corydiidae | B081 | Breeding colony of Kyle Kandilian | N/A | N/A |
| ***Eurycotis decipiens*** | Blattidae | B071 | Breeding colony of Kyle Kandilian | N/A | N/A |
| ***Galiblatta cribrosa*** | Blaberidae | Z98 | Nouragues, French Guiana | N/A | 14-Jun-2015 |
| ***Gromphadorhina grandidieri*** | Blaberidae | B030 | Breeding colony of Kyle Kandilian | N/A | N/A |
| ***Gyna capucina*** | Blaberidae | Z139GY | Ebogo, Cameroon | Frantisek Juna | 08-Sep-2015 |
| ***Ischnoptera deropeltiformis*** | Ectobiidae | B083 | Torreya State Park, Bristol, Florida, USA | | 07-Jul-2009 |
| ***Lamproblatta* sp.** | Lamproblattidae | LA male | Petit Saut, French Guiana | Frantisek Juna | 08-Jul-2009 |
| ***Laxta* sp.** | Blaberidae | AUS2 | Olney State Forest, New South, Wales, Australia | Nathan Lo and Thomas Bourguignon | 25-Aug-2015 |
| ***Macropanesthia rhinoceros*** | Blaberidae | B092 | Breeding colony of Kyle Kandilian | N/A | N/A |
| ***Mastotermes darwiniensis*** | Isoptera | MADAR | GenBank | N/A | N/A |
| ***Megaloblatta* sp.** | Ectobiidae | ECMD1 | Podocarpus National Park, Ecuador | Frantisek Juna | Apr-2016 |
| ***Melanozosteria* sp.** | Blattidae | Melanozosteria_sp. | Cairns, Queensland, Australia | David Rentz | 18-Dec-2015 |
| ***Methana* sp.** | Blattidae | AUS1 | North Manly, New South Wales, Australia | Nathan Lo | 01-Aug-2015 |

| | | | | | |
|---|---|---|---|---|---|
| ***Nauphoeta cinerea*** | Blaberidae | BNCIN | GenBank | N/A | N/A |
| ***Neolaxta mackerrasae*** | Blaberidae | B107 | Paluma Range, Queensland, Australia | David Rentz | 15-Oct-2015 |
| ***Opisthoplatia orientalis*** | Blaberidae | Z15100 | Breeding colony of J. Hromádka | N/A | N/A |
| ***Panchlora nivea*** | Blaberidae | B044 | Breeding colony of Kyle Kandilian | N/A | N/A |
| ***Panesthia angustipennis*** | Blaberidae | Z138 | Breeding colony in Czech Republic, orig. Vietnam | N/A | 01-Aug-2015 |
| ***Panesthia* sp.** | Blaberidae | Panesthia_sp | Bubeng, Yunnan, China | N/A | 08-Jul-2009 |
| ***Paranauphoeta circumdata*** | Blaberidae | PARA | N/A | N/A | N/A |
| ***Paratemnopteryx couloniana*** | Ectobiidae | B061 | Breeding colony of Kyle Kandilian | N/A | N/A |
| ***Parcoblatta virginica*** | Ectobiidae | B102 | Breeding colony of Kyle Kandilian | N/A | N/A |
| ***Periplaneta americana*** | Blattidae | BPLAN | GenBank | N/A | N/A |
| ***Phyllodromica* sp.** | Ectobiidae | Phil | Czech Republic | Thomas Bourguignon | 01-Aug-2015 |
| ***Platyzosteria* sp.** | Blattidae | AUS3 | Olney State Forest, New South, Wales, Australia | Nathan Lo and Thomas Bourguignon | 25-Aug-2015 |
| ***Protagonista lugubris*** | Blattidae | Cockroach contig 4907 | Mt Diaoluo, Hainan, China | Zongqing Wang | 25-May-2015 |
| ***Pycnoscelus femapterus*** | Blaberidae | B048 | Breeding colony of Kyle Kandilian | N/A | N/A |
| ***Rhabdoblatta* sp.** | Blaberidae | RHA | Kuranda, Queensland, Australia | David Rentz | 16-Sep-2015 |
| ***Shelfordella lateralis*** | Blattidae | B080 | Breeding colony of Kyle Kandilian | N/A | N/A |
| ***Therea regularis*** | Corydiidae | B091 | Palm plantation between Puducherry and Auroville, India | Kyle Kandilian | N/A |
| ***Tryonicus parvus*** | Tryonicidae | Tryonicus_parvus | Olney State Forest, New South, Wales, Australia | Nathan Lo and Thomas Bourguignon | 10-Mar-2016 |
| ***Sulcia muelleri*** | Flavobacteriaceae | CARI | GenBank | N/A | N/A |
| ***Sulcia muelleri*** | Flavobacteriaceae | PSPU | GenBank | N/A | N/A |
| ***Sulcia muelleri*** | Flavobacteriaceae | CARI | GenBank | N/A | N/A |
| ***Flavobacterium gilvum*** | Flavobacteriaceae | EM1308 | GenBank | N/A | N/A |
| ***Lutibacter* sp.** | Flavobacteriaceae | LPB0138 | GenBank | N/A | N/A |
| ***Tenacibaculum dicentrarchi*** | Flavobacteriaceae | AY7486TD | GenBank | N/A | N/A |
| ***Polaribacter* sp.** | Flavobacteriaceae | LT629752 | GenBank | N/A | N/A |

**Supplementary table S2.** A list of GenBank accession numbers and names of all 104 *Blattabacterium* genes used for this study.

| Accession No. | Gene name |
|---|---|
| **MN038417 - MN038462** | Dihydrolipoyllysine-residue succinyltransferase component of 2oxoglutarate dehydrogenase complex. |
| **MN038463 - MN038510** | Cysteine desulfuration protein |
| **MN038511 - MN038558** | hypothetical protein |
| **MN038559 - MN038606** | Putative 1,2-phenylacetyl-CoA epoxidase, subunit D |
| **MN038607 - MN038654** | UDP-N-acetylglucosamine--N-acetylmuramyl-pentapeptide pyrophosphoryl-undecaprenol N-acetylglucosamine transferase |
| **MN038655 - MN038703** | 50S ribosomal protein L11 |
| **MN038704 - MN038752** | 50S ribosomal protein L11 |
| **MN038753 - MN038800** | Fumarate reductase flavoprotein subunit |
| **MN038801 - MN038848** | Glutamate dehydrogenase |
| **MN038849 - MN038895** | Asparagine tRNA ligase |
| **MN038896 - MN038942** | Polyribonucleotide nucleotidyltransferase |
| **MN038943 - MN038989** | RNA polymerase sigma factor SigA |
| **MN038990 - MN039036** | 3-oxoacyl-acyl-carrier-protein synthase 2 |
| **MN039037 - MN039083** | Acyl carrier protein |
| **MN039084 - MN039130** | ATP synthase epsilon chain |
| **MN039131 - MN039177** | 30S ribosomal protein S2 |
| **MN039178 - MN039223** | 50S ribosomal protein L13 |
| **MN039224 - MN039271** | 10 kDa chaperonin |
| **MN039272 - MN039318** | 60 kDa chaperonin |
| **MN039319 - MN039365** | Glutamine tRNA ligase |
| **MN039366 - MN039412** | DNA-directed RNA polymerase subunit beta' |
| **MN039413 - MN039459** | Glyceraldehyde-3-phosphate dehydrogenase A |

| | |
|---|---|
| **MN039460 - MN039506** | 50S ribosomal protein L21 |
| **MN039507 - MN039553** | 50S ribosomal protein L22 |
| **MN039554 - MN039600** | 30S ribosomal protein S19 |
| **MN039601 - MN039647** | 50S ribosomal protein L2 |
| **MN039648 - MN039695** | 50S ribosomal protein L3 |
| **MN039696 - MN039742** | 50S ribosomal protein L1 |
| **MN039743 - MN039790** | Cysteine desulfurase SufS |
| **MN039791 - MN039837** | FeS cluster assembly protein SufB |
| **MN039838 - MN039884** | Protein translocase subunit SecY |
| **MN039885 - MN039931** | 50S ribosomal protein L15 |
| **MN039932 - MN039979** | 30S ribosomal protein S10 |
| **MN039980 - MN040027** | Elongation factor G |
| **MN040028 - MN040075** | 30S ribosomal protein S7 |
| **MN040076 - MN040123** | 30S ribosomal protein S12 |
| **MN040124 - MN040171** | Methionine aminopeptidase 1 |
| **MN040172 - MN040219** | 30S ribosomal protein S5 |
| **MN040220 - MN040267** | Alternate 30S ribosomal protein S14 |
| **MN040268 - MN040314** | 50S ribosomal protein L14 |
| **MN040315 - MN040361** | 30S ribosomal protein S17 |
| **MN040362 - MN040408** | 50S ribosomal protein L16 |
| **MN040409 - MN040456** | 30S ribosomal protein S3 |
| **MN040457 - MN040504** | Two names: 1)] Acetylornithine deacetylase; 2)] Succinyl-diaminopimelate desuccinylase |
| **MN040505 - MN040552** | Aspartate semialdehyde dehydrogenase |
| **MN040553 - MN040599** | 50S ribosomal protein L17 |

| | |
|---|---|
| **MN040600 - MN040647** | DNA-directed RNA polymerase subunit alpha |
| **MN040648 - MN040695** | 30S ribosomal protein S4 |
| **MN040696 - MN040743** | 30S ribosomal protein S11 |
| **MN040744 - MN040791** | 30S ribosomal protein S13 |
| **MN040792 - MN040839** | Translation initiation factor IF-1 |
| **MN040840 - MN040887** | N-acetylornithine carbamoyltransferase |
| **MN040888 - MN040935** | Carbamoyl-phosphate synthate large chain |
| **MN040936 - MN040983** | Carbamoyl-phosphate synthate small chain |
| **MN040984 - MN041031** | Acetylornithine aminotransferase |
| **MN041032 - MN041079** | N-acetyl-gamma-glutanyl-phosphate reductase |
| **MN041080 - MN041127** | Argininosuccinate synthase |
| **MN041128 - MN041175** | 30S ribosomal protein S1 |
| **MN041176 - MN041223** | Enoyl-acyl-carrier-protein reductase NADH FabI |
| **MN041224 - MN041271** | S-adenosylmethionine synthase |
| **MN041272 - MN041319** | Phospho-2-dehydro-3-deoxyheptonate aldolase |
| **MN041320 - MN041366** | 50S ribosomal protein L9 |
| **MN041367 - MN041412** | 30S ribosomal protein S6 |
| **MN041413 - MN041460** | tRNA modification GTPase MnmE |
| **MN041461 - MN041508** | Lon protease 2 |
| **MN041509 - MN041556** | Histidine--tRNA ligase |
| **MN041557 - MN041603** | Phenylalanine--tRNA ligase alpha subunit |
| **MN041604 - MN041650** | DNA gyrase subunit B |
| **MN041651 - MN041698** | 30S ribosomal protein S16 |
| **MN041699 - MN041746** | Aspartate aminotransferase |

| | |
|---|---|
| **MN041747 - MN041794** | Lysine--tRNA ligase |
| **MN041795 - MN041840** | Octanoyltransferase |
| **MN041841 - MN041887** | Methionine--tRNA ligase |
| **MN041888 - MN041935** | Histidinol dehydrogenase |
| **MN041936 - MN041981** | hypothetical protein |
| **MN041982 - MN042029** | Phosphate acetyltransferase |
| **MN042030 - MN042076** | 1-deoxy-D-xylulose-5-phosphate synthase |
| **MN042077 - MN042124** | Transketolase 2 |
| **MN042125 - MN042170** | SsrA-binding protein |
| **MN042171 - MN042216** | Lipoyl synthase |
| **MN042217 - MN042264** | Multifunctional CCA protein |
| **MN042265 - MN042311** | 2,3,4,5-tetrahydropyridine-2,6-dicarboxylate N-succinyltransferase |
| **MN042312 - MN042358** | putative branched-chain-amino-acid aminotransferase |
| **MN042359 - MN042405** | 2-oxoisovalerate dehydrogenase subunit beta |
| **MN042406 - MN042452** | Ribosomal RNA small subunit methyltransferase A |
| **MN042453 - MN042499** | Putative aminopeptase YsdC |
| **MN042500 - MN042546** | Diaminopimelate epimerase |
| **MN042547 - MN042594** | ATP synthase subunit c |
| **MN042595 - MN042641** | ATP synthase subunit beta |
| **MN042642 - MN042688** | Chaperone protein DnaJ |
| **MN042689 - MN042735** | hypothetical protein |
| **MN042736 - MN042782** | Ribose-phosphate pyrophosphokinase |
| **MN042783 - MN042830** | Imidazole glycerol phosphate synthase subunit HisF |
| **MN042831 - MN042878** | Imidazole glycerol phosphate synthase subunit HisH |

| | |
|---|---|
| **MN042879 - MN042926** | Ribosome recycling factor |
| **MN042927 - MN042973** | ATP synthase subunit A |
| **MN042974 - MN043021** | Bifunctional aspartokinase |
| **MN043022 - MN043069** | hypothetical protein |
| **MN043070 - MN043116** | 3-dehydroquinate synthase |
| **MN043117 - MN043164** | DNA gyrase subunit A |
| **MN043165 - MN043211** | Fructose-bisphosphate aldolase |
| **MN043212 - MN043259** | hypothetical protein |
| **MN075834-MN075880** | Elongation Factor |
| **MN075881-MN075928** | tRNA-2-methylthio-N6-dimethylallyladenosine synthase |
| **CP003535-CP003536** | *Blaberus giganteus* |
| **CP003605-CP003606** | *Blatta orientalis* |
| **CP001487** | *Blattella germanica* |
| **CP003015-CP003016** | *Cryptocercus punctulatus* |
| **CP003000, CP003095** | *Mastotermes darwiniensis* |
| **CP005488-CP005489** | *Nauphoeta cinerea* |
| **CP001429-CP001430** | *Periplaneta americana* |
| **CP002163** | *Sulcia muelleri* |
| **AP013293** | *Sulcia muelleri* |
| **CP002165** | *Sulcia muelleri* |
| **CP017479** | *Flavobacterium gilvum* |
| **CP017478** | *Lutibacter* sp. |
| **CP013671** | *Tenacibaculum dicentrachi* |
| **KT25b** | *Polaribacter* sp. |

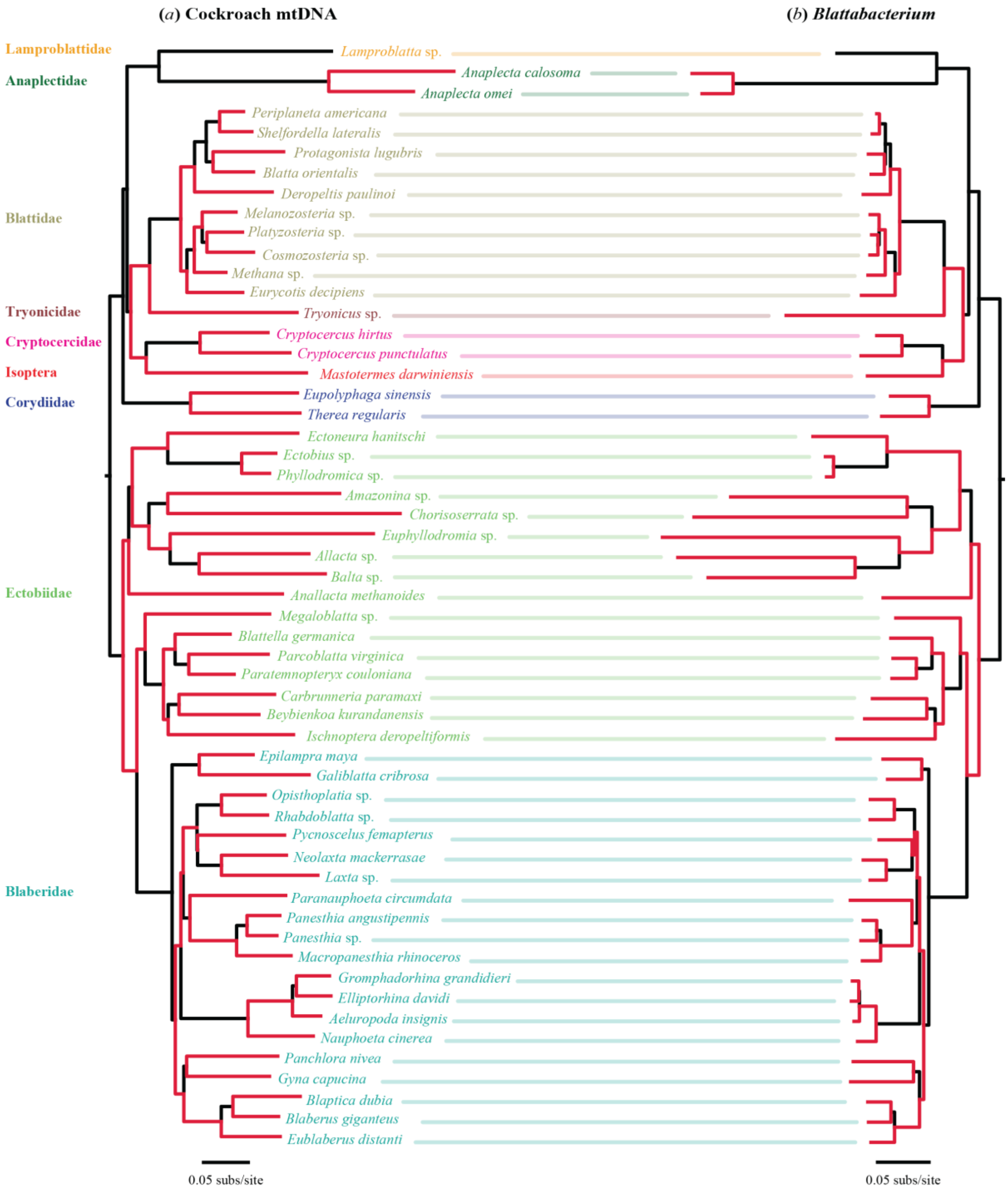


**Supplementary Figure S1**. Phylogenetic trees of (*a*) cockroaches and (*b*) their *Blattabacterium* symbionts inferred using maximum likelihood in RAxML. A fixed topology (obtained from the *Blattabacterium* tree shown in figure 1) was used in each analysis. Twenty-seven phylogenetically independent pairs of lineages used to test for correlations of evolutionary rates are shown in red. Species names are coloured according to the family to which they belong, as shown on the left of the figure.

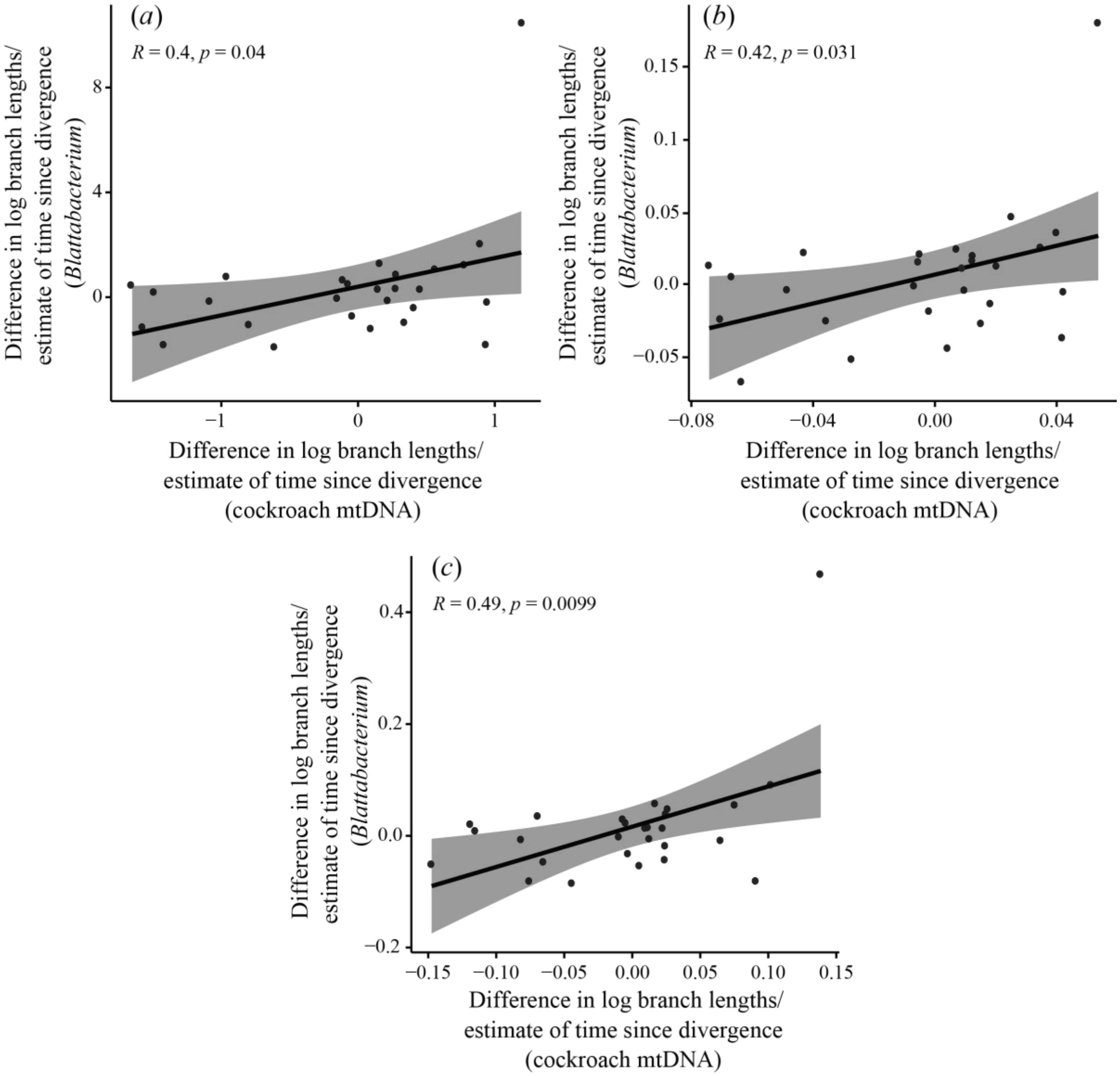


**Supplementary Figure S2.** Standardized tests for correlation of molecular evolutionary rates between 27 independent pairs of *Blattabacterium* and host cockroach mitochondria, based on protein-coding genes. Three standardizations were carried out, each based on dividing log-transformed branch-length differences by the square root of an estimate of time since divergence for the pair. In the first standardization (*a*), time since divergence for host pairs was estimated as the average branch length of the host pair, divided by an assumed rate of 0.001 subs/site/million years, while for corresponding symbionts it was estimated as the average branch length of the symbiont pair, divided by the same assumed rate. In the second (*b*) and third (*c*) standardizations, times since divergence for both symbionts and hosts were based either on average branch lengths of host pairs only or symbiont pairs only.

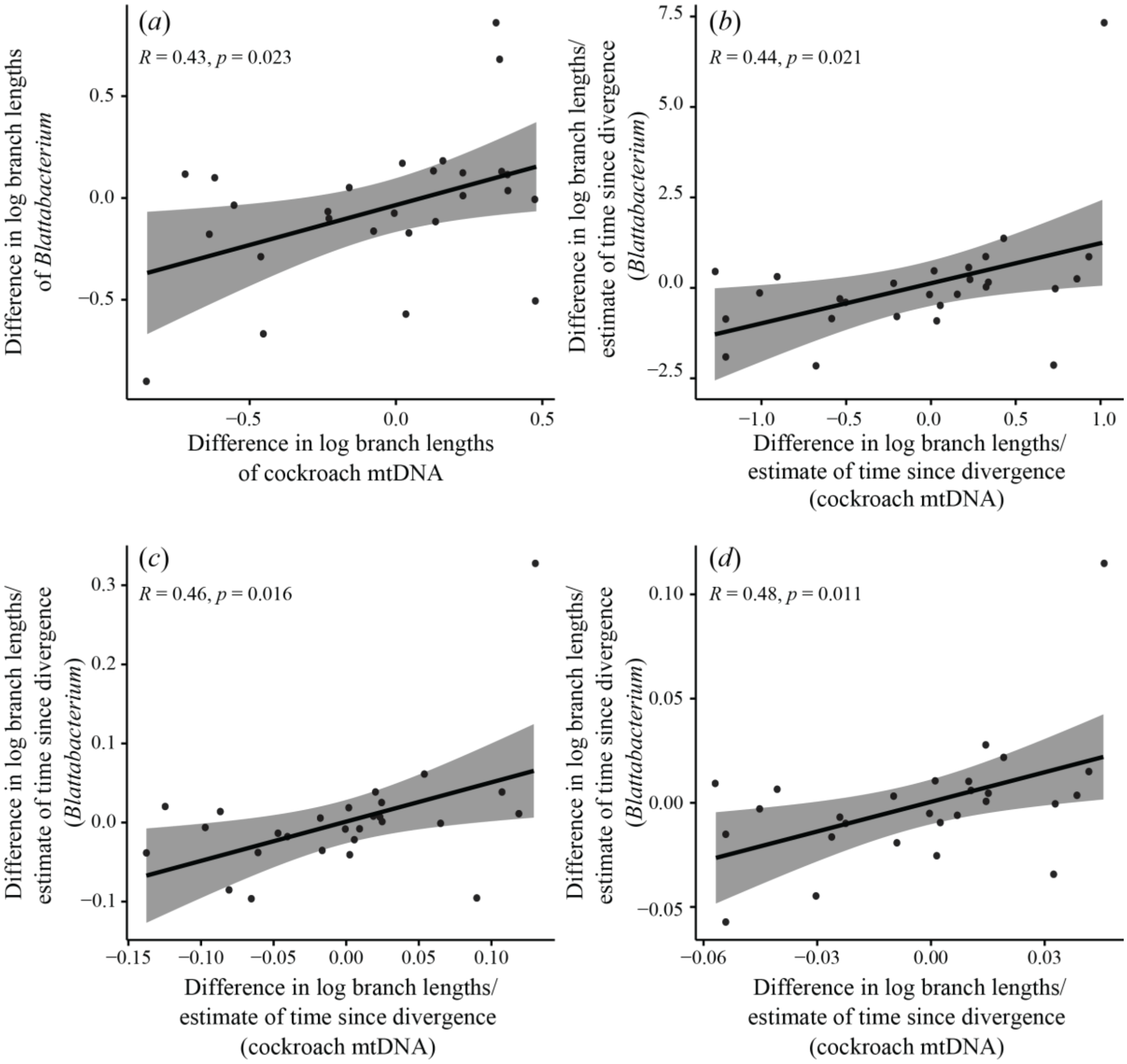


**Supplementary Figure S3**. Tests for correlation of molecular evolutionary rates between 27 independent pairs of *Blattabacterium* and host cockroach mitochondria, based on amino acid data translated from protein-coding genes. (*a*) Test based on comparison of log-transformed branch-length differences. Three standardizations were also carried out, each based on dividing log-transformed branch-length differences by the square root of an estimate of time since divergence for the pair. In the first standardization (*b*), time since divergence for host pairs was estimated as the average branch length of the host pair, divided by an assumed rate of 0.001 subs/site/million years, while for corresponding symbionts it was estimated as the average branch length of the symbiont pair, divided by the same assumed rate. In the second (*c*) and third (*d*) standardizations, times since divergence for both symbionts and hosts were based either on average branch lengths of host pairs only or symbiont pairs only.

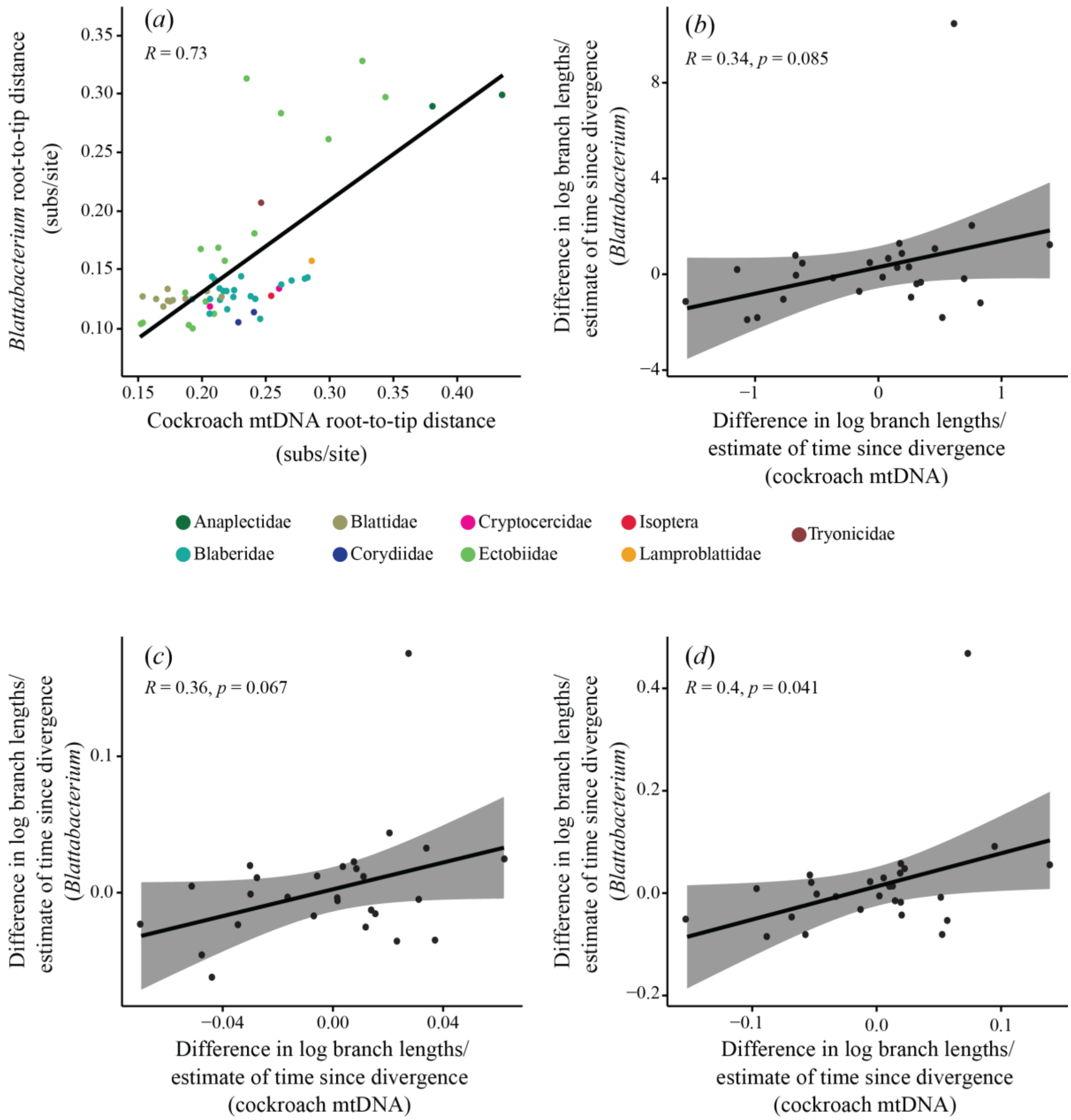


**Supplementary Figure S4.** Comparison of evolutionary rates of *Blattabacterium* symbionts and their host cockroaches, based on protein-coding genes from host and symbiont, plus rRNAs+tRNAs from host mitochondria. (*a*) Correlation of root-to-tip distances in phylogenies of *Blattabacterium* and cockroaches. (*b*–*d*) Standardized tests for correlation of molecular evolutionary rates between 27 independent pairs of *Blattabacterium* and host cockroach mitochondria. Three standardizations were carried out, each based on dividing log-transformed branch-length differences by the square root of an estimate of time since divergence for the pair. In the first (*b*), time since divergence for host pairs was estimated as the average branch length of the host pair, divided by an assumed rate of 0.001

subs/site/million years, while for corresponding symbionts it was estimated as the average branch length of the symbiont pair, divided by the same assumed rate. In the second (*c*) and third (*d*) standardizations, times since divergence for both symbionts and hosts were based either on average branch lengths of host pairs only or symbiont pairs only.
